# Visual and vestibular processing of vertical motion: a psychophysical study

**DOI:** 10.1101/2024.12.15.628168

**Authors:** Sergio Delle Monache, Barbara La Scaleia, Anna Finazzi Agrò, Francesco Lacquaniti, Myrka Zago

## Abstract

The motion of objects and ourselves along the vertical is affected by gravitational acceleration. However, the visual system is poorly sensitive to accelerations, and the vestibular otoliths do not disassociate gravitational and inertial accelerations of ego-motion. Here, we tested the hypothesis that the brain resolves visual and vestibular ambiguities about vertical motion with internal models of gravity, which predict that downward motions are accelerated and upward motions are decelerated by gravity. In visual sessions, a target moved up or down while participants remained stationary. In vestibular sessions, participants were moved up or down, while they fixated an imaginary target moving along. In visual-vestibular sessions, participants were moved up or down while the visual target remained fixed. We found that downward motions of either the visual target or the participant were systematically perceived as lasting less than upward motions of the same duration, and vice-versa for the opposite direction of motion, consistent with the prior assumption that downward motion is accelerated and upward motion is decelerated by gravity. In visual-vestibular sessions, there was no significant difference in the average estimates of duration of downward and upward motion of the participant. However, there was large inter-subject variability of these estimates.

## INTRODUCTION

We move and experience object motion in a 3D-environment. Our spatial perception and orientation in 3D are anchored to the gravitational vertical and the horizon, and depend heavily on visual and vestibular cues. Thus, in the natural visual scenes commonly experienced, there is as much image structure at vertical orientation as at horizontal orientation, and least at obliques^1^. With regards to the vestibular inputs experienced in daily life, small-amplitude self-motion in the vertical direction is as common as that in the horizontal direction^2^. Neural processing of object motion and self-motion in the horizontal and vertical directions is partially segregated in the brain^3,4^. Here, we are concerned with the perception of motion of objects or ourselves in the vertical direction.

In order to process the motion of visual targets or our own motion with a vertical component, we must take gravitational acceleration (1*g*) into account^5,6^. However, the visual system is poorly sensitive to accelerations^7^, while the otolith organs of the vestibular system cannot disentangle gravitational and inertial accelerations of self-motion^8^. Therefore, it has been hypothesized that the brain accounts for gravity by complementing multisensory information with internal models of gravity effects^9–11^. These internal models consist of neural mechanisms mimicking the kinematics under gravity in an approximate, probabilistic manner^12–18^. Neural responses coherent with these internal models have been described in visual-vestibular brain regions with electrophysiological recordings in monkeys^19,20^, and functional magnetic resonance imaging (fMRI) and transcranial magnetic stimulation (TMS) in humans^3,21–25^.

Behavioural evidence for the motor utilization of a gravity model has been provided by showing that humans can catch falling targets in the absence of visual cues when interception time is predictable^26,27^. In the extrapolation of vertical target motion through a visual occlusion, the interception performance of targets accelerated under gravity (1*g*) is always superior to that of targets decelerated under reversed gravity (-1*g*), or moving at constant speed (0*g*)^28^. Moreover, the implicit, erroneous assumption of 1*g* leads astronauts to move too early to catch a ball descending at constant speed (0*g*) in weightlessness^29^, as it does for participants asked to punch a virtual target descending at 0*g* on Earth^30^. A different study showed a response bias as a function of the upward or downward direction of target motion: subjects triggered movements to intercept a virtual target earlier when the target came from above instead of below, consistent with the prior assumption that downward motion is accelerated by gravity while upward motion is decelerated by gravity^31^. It has also been shown that the response bias for accelerated 1*g* motion in the upward direction (unnatural motion) versus the downward direction (natural motion) depends on the familiarity with the target trajectory^32^. Vestibular inputs also play a role in interception timing^33^, since this response bias reversed sign between the above and below conditions, in parallel with the sign reversal of otolith signals at the transition from hypergravity to hypogravity during parabolic flight^34^. Eye movements also betray the implementation of a gravity model, as shown by the higher gain of smooth pursuit when tracking targets moving at 1*g* than those moving at 0*g* or other *g* levels^35–37^, and by faster and smoother pursuit in response to downward versus upward motion^38^. As for the vestibular system, reflexive eye movements in response to tilts and translations appear related to central estimates of linear acceleration derived from an internal model of gravity, rather than related to the net gravito-inertial force encoded by the otoliths^39–41^.

Perceptual responses also appear compatible with internal models of gravity^42^. Thus, the precision of visual motion duration is significantly better for motion accelerating downwards than accelerating upwards^43,44^. The threshold (precision) of self-motion vestibular discrimination in the dark has also been reported to be significantly better for motion accelerating downwards than accelerating upwards in some studies^45–48^, while no significant difference has been reported in other studies^49^. Visual motion coherence thresholds are significantly lower when the observer’s position and target motion are congruent with gravity^50^. A downward bias also exists in the ability to detect visual accelerations^51^. Visual preference for ecological downward accelerating motion arises early in infancy^52^.

The best evidence for an internal model, as opposed to direct sensory measures, is offered by misperceptions or biased perceptions^53^. According to von Helmholtz (1867)^54^, ‘It is just those cases that are not in accordance with reality which are particularly instructive for discovering the laws of the processes by which normal perception originates’. Thus, Moscatelli et al. (2019)^55^ showed that downward visual motions at constant speed were misperceived as faster than equally-lasting upward motions. Relatedly, Miwa et al. (2019)^56^ found that observers correctly identify as uniform an upward motion at constant speed, but they erroneously judge downward accelerating motion as uniform (see also La Scaleia et al. 2014^57^ and Phan et al. 2024^58^). Indeed, observers detect the presence of an acceleration phase in targets moving upwards more easily than in targets moving downwards^59^. In a visual search task, observers find targets undergoing acceleration/deceleration (at 1*g*) among distractors moving at constant speed (0*g*) more easily when all objects move upwards or horizontally than when they move downwards^60^. Moreover, previously viewed descending targets are judged as being displaced forward along the path more than do ascending targets, consistent with a representational gravity^61^.

Biases may also arise in the vestibular perception of self-motion due to asymmetric labyrinthine information or asymmetric central processing of peripheral information^62^. Thus, the horizontal component of motion relative to the vertical component is overestimated during two-dimensional motions in the sagittal (elevation) plane^63^. This bias could reflect the higher sensitivity of the utricles relative to the saccules^8^. However, clear-cut biases in the up-down direction similar to those reported for the visual stimuli have not been reported so far for vestibular perception of self-motion^49,63,64^. By comparing direction discrimination for naso-occipital, inter-aural, cranio-caudal motions in upright, supine and side-lying subjects, Kobel et al. (2021)^47^ found no significant bias in any of the individual conditions, but a significant main effect of motion relative to gravity due to a larger positive bias in earth-vertical motions than earth-horizontal motions for 2-Hz displacements. Because no impact of gravity on vestibular bias was found for 1-Hz displacements where vestibular contributions are known to dominate^65^, Kobel et al. interpreted the bias at 2-Hz as resulting from up-down asymmetries in somesthetic cues. A similar argument was put forth by Nesti et al. (2014)^46^. However, both Nesti’s and Kobel’s studies were mainly focused on the assessment of vestibular perceptual precision (thresholds) rather than the assessment of accuracy (bias).

Here, we searched for perceptual directional biases in the discrimination of vertical motion duration. Participants sat upright on a motion platform and wore a virtual-reality headset. In visual sessions (VI), a visual target moved up or down while the participant remained stationary. In vestibular sessions (VE), the participant was moved up or down and was asked to fixate an imaginary target moving together with the subject. In visual-vestibular sessions (VV), the participant was moved up or down while the visual target was displayed at a fixed location, thus shifting relative to the subject in the direction opposite to their movement. The acceleration profile of the motion consisted of one cycle of a sinewave. In a two-alternative forced choice task, participants were asked to judge which one of two stimuli -presented consecutively in random, balanced order-had the shorter duration, i.e., a comparison stimulus moving up or down and a reference stimulus moving in the opposite direction (down or up). While the duration of the reference was fixed at 1.400 s, the duration of the comparison varied randomly between 0.840 s and 1.960 s. Thus, all stimuli were at a relatively low frequency (between about 0.5 Hz and 1.2 Hz). In VE sessions, these frequencies corresponded to those thought to maximize vestibular over somesthetic contributions during self-motion perception^65,66^.

We computed the point of subjective equality (PSE) of the psychometric functions, i.e. the value of a comparison stimulus in one direction that is equally likely to be judged as lasting more or less than that of the reference stimulus in the opposite direction. The PSE reflects the presence of possible perceptual biases^67^. The hypothesis of an internal model of gravity predicts that the downward motion of the target in the visual (VI) sessions and that of the participant in the vestibular (VE) sessions should be perceived as lasting significantly less (positive bias or greater PSE) than the upward motions of the same duration, and vice-versa for the opposite direction of motion. This prediction would be consistent with the prior assumption that downward motion is accelerated by gravity while upward motion is decelerated by gravity^31^.

The prediction for the visual-vestibular sessions (VV), however, is not univocal due to the potential ambiguity in the subjective interpretation of the stimuli, given that the participant was moved up or down while the visual target was displayed at a fixed location. Therefore, the target shifted relative to the participant in the opposite direction to the participant motion. If the participant responded mainly to either the vestibular or the visual information, we should find a bias similar to that of the corresponding VE session or the VI session, respectively. If, however, the participant responded to both the vestibular and the visual cues at the same time, the visual and the vestibular bias should roughly cancel out, given that the resulting retinal slip would tend to compensate the head motion.

## METHODS

### Participants

Twenty young adults (10 women, 10 men, mean age 29.7 ± 6.0 SD), with normal or corrected-to-normal vision, no history of psychiatric, neurological or vestibular symptoms, dizziness or vertigo, motion-sickness susceptibility, major health problems or medications potentially affecting vestibular function, volunteered to participate in the experiments. Sample size was calculated to detect an effect size of 0.83 (Cohen’s d, estimated from previous studies with comparable conditions – Moscatelli et al.^55^ and Kobel et al.^47^ – as well as from our preliminary data) with power of 0.8 and alpha of 0.017 (i.e. 0.050 / 3, where 3 is the correction for the expected levels of comparison corresponding to the number of sensorial modalities involved. G*power version 3.1). Data were anonymized after collection for subsequent analysis. To avoid experimenter-expectancy effects, the analysis was carried out on blinded data by experimenters different from those involved in data collection. All participants gave written informed consent to procedures approved by the Institutional Review Board of Santa Lucia Foundation (protocol n. CE/PROG0.757), in conformity with the Declaration of Helsinki (World Medical Association) regarding the use of human participants in research.

### Setup

Participants sat on a gaming chair placed on a six-degrees-of-freedom motion platform (MB-E-6DOF/12/1000, Moog, USA). A 4-point harness held their trunk securely in place, while a medium density foam pad under the feet minimized plantar cues about body displacement. For acoustic isolation, participants wore earplugs (1100 series, 3M, USA) and on-ear headphones (Anker Soundcore Q20, China). They wore a 3D VR-headset system (Quest Pro, Meta, USA) with a refresh rate of 90 Hz and a horizontal Field of View of 106°. The VR-headset system also acquired the position of the head and eyes at 90 Hz, the latter with an average accuracy better than 2 degrees^68,69^. Data from platform (chair) and VR-headset (head and eyes position and orientation) were referred to a fixed world reference system (W_RF_, see Figure 1). We estimated the roto-translation matrix required to map the 3D data acquired in the VR-headset (head) reference frame and in the platform reference frame into W_RF_ by performing a spatial calibration procedure before each experimental session with the VR-headset mounted on the platform in a stable position. All details about the general spatial calibration procedures can be found in La Scaleia et al.^70^. The temporal alignment of feedback positions and orientations of the VR-headset (head) and of the motion platform (chair) was obtained by means of a cross-correlation analysis. We found a lag of about 40 ms, which was then taken into account. The data relative to the position of the chair, head and eyes acquired during the experiment and referred to W_RF_ were used to assess that the participants kept the head and eyes roughly aligned with the straight-ahead direction (see below) during the execution of the task.

**Figure 1.**
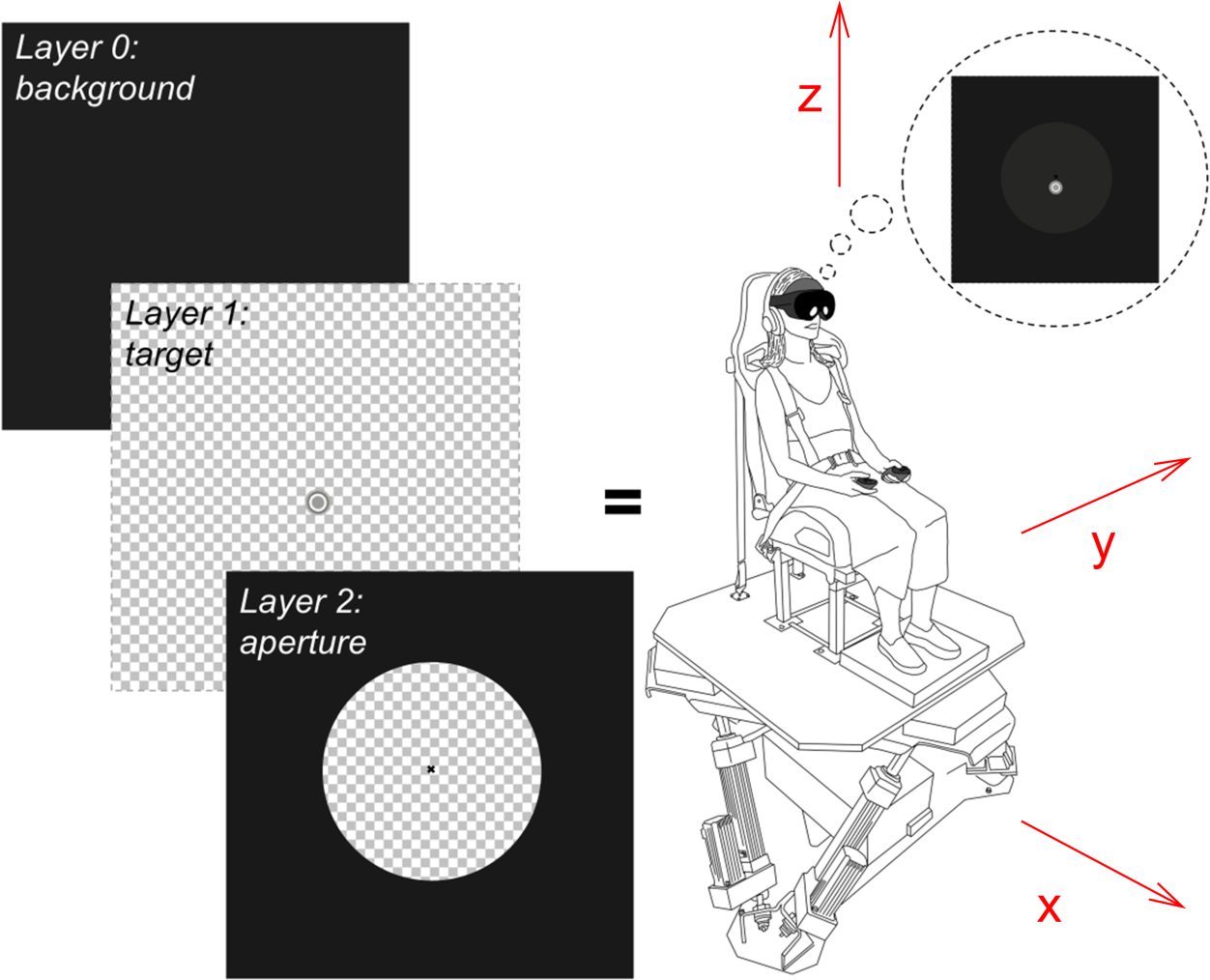
Visual stimulation in the visual and visuo-vestibular sessions consisted of three superimposed layers, oriented perpendicular to the subject’s antero-posterior axis from back to front (see Methods). Layer 0 comprised the background of the virtual scene. Layer 1, positioned immediately in front of Layer 0, contained the visual target, a grey-textured disk with a diameter of 1.68° of visual angle. Located in front of Layer 1, Layer 2 included two elements: a uniformly colored surface with a central circular aperture; and a black cross tilted by 45° at its center. In Layer 1 and Layer 2, transparent portions of the scenario are represented with a squared texture. The resulting 3D virtual scenario was perceived by the subject through the VR-headset system (see also Supplementary Video 1 and Supplementary Video 3). In the vestibular session, visual stimulation consisted solely of the background (Layer 0; see also Supplementary Video 2). The W_RF_ (axes x, y, z, represented in red in the figure) has its origin in the flying base motion centroid position when the flying base is in its home settled position. Motion centroid is the centroid of the joints below the flying base of the motion platform. The z-axis is orthogonal to the ground and oriented upwards. The x-axis is oriented along the posterior-anterior direction of the chair.

### Scenario

With the VR-headset system, participants were presented with a three-dimensional scenario programmed by means of Unity engine (editor version 2021.3.25f1, packed with XR Plugin Management 4.3.3, Oculus XR Plugin 3.3.0 and Meta Movement SDK 1.4.1). Prior to the beginning of the experiment, each subject was asked to slide the lenses of the VR-headset closer or further apart from one another so as to have a good VR experience and to achieve the best image clarity. This was required to take into account the interpupillary distance of the participant. To this end, they had to fuse binocularly a text presented in stereo at the same distance as the experimental scene. Afterwards, participants underwent the eye-tracking calibration routine included in the VR-headset software.

Each experimental session started with the same initial calibration scene, consisting of a plain, uniform scene (the intensity of each color in the RGB color code was set to 20). Participants were asked to keep their head in a relaxed position, facing forward along the perceived antero-posterior direction and looking straight-ahead. The mean position of the eyes acquired over 1 s during the calibration phase was used to define the position of the working area. This area was placed 42 cm in front of the participant, along their sightline, and was roughly perpendicular to the world reference ground. Afterwards, the location of the working area was held constant in world reference, independently of the subject’s actual position and orientation. By contrast, the point of view was updated according to the line of sight estimated through the convergence of the gaze directions of the two eyes recorded by the VR-headset at 90 Hz throughout the experimental session.

The working area consisted of a group of virtual objects organized in three overlayed layers (Figure 1). These three layers were oriented roughly perpendicularly to the world reference ground (plane x-y, see W_RF_ in Figure 1 for further details), with their surfaces exposed to the subject. They were illuminated by an orthogonal directional white light (RGB color intensity set to 255 for each color). Layer 0, the background layer, was a uniformly grey surface (see below for the RGB color intensity). In front of this, Layer 1 contained the only visual target in a transparent environment (represented with a squared texture in Figure 1). The target was a disk with a diameter of 1.68° of visual angle, featuring a grey texture with high contrast against the Layer 0 (Michelson contrast 74%). This grey texture consisted of four concentric circles, each displaying a distinct grayscale color level (RGB color intensity set to 136, 89, 204, 102 for each color, from outer to inner circles, respectively), similarly to the high-contrast condition of Moscatelli et al.^55^. Further anteriorly, Layer 2 included two elements: a black (RGB color intensity set to 0 for each color) fixation cross tilted by 45° measuring 0.42° of visual angle in both width and height, and a uniformly colored vertical surface (RGB color intensity set to 25 for each color) with a circular transparent aperture (in Figure 1, the transparency was represented with the same squared texture as for Layer 1) centered on the fixation cross, with a diameter of 14.25° of visual angle. These three layers were superimposed onto each other, and were oriented roughly perpendicularly to the world reference ground, with their surfaces exposed to the subject.

The experiment consisted of three sessions, each involving vertical movements of different elements of the virtual scenario and/or of the chair. (i) In the visual session (VI), all Layers of the visual scene were visible, the chair remained static throughout the session, as did the Layer 0 (background; RGB color intensity set to 30 for each color) and the Layer 2. In its initial position, the target in Layer 1 was either vertically above or vertically below the fixation cross of Layer 2. The target became visible through the aperture when the Layer 1 moved vertically (see Supplementary Video 1). When the target, during Layer 1 motion, was aligned with the fixation cross of Layer 2, it was partially hidden by the cross (Layer 2 was stacked on top of Layer 1). (ii) In the vestibular session (VE), only the Layer 0 (background) was visible in the virtual scene (Layer 1 and Layer 2 were hidden from view) while the chair moved vertically (see Supplementary Video 2). To match the overall luminosity of the virtual scenario between sessions, in VE the Layer 0 was slightly darker than in the other sessions (RGB color intensity set to 27 for each color), (iii) In the visuo-vestibular session (VV), the chair moved vertically as in the VE session. All three layers of the visual scene were visible as in VI. Unlike VI, the visual target in Layer 1 was stationary, immediately above or below the aperture of Layer 2, however Layer 2 position was linked to that of the chair movement. As the chair moved, so did the aperture and the fixation cross of Layer 2, making the target temporarily visible during the stimulation (see Supplementary Video 3). In the VV session, the positions of the chair and Layer 2 (aperture and fixation cross) were synchronized by adjusting the timing between the platform motion and VR-headset updates. To assess the lag between the two systems, we measured the onset of platform motion by means of a MEMS triaxial accelerometer (MPU-6050, InvenSense, US), and the onset of visual motion by means of a photodiode (BPW21 Siemens, Germany). We found that platform motion lagged behind visual motion by ∼40 ms. To compensate for this lag, we shifted the first update of the position of Layer 2 by 40 ms in each trial, so that the onset of visual motion was synchronized with the start of platform motion. The position of Layer 2 was then updated at 90 Hz in a feedforward manner following the same law of motion as the chair.

### Stimuli

The path of the stimuli was identical across conditions and sessions (VI, VE and VV), but the direction was either downward or upward. The length of the path (Δp) of either the visual target or the chair was 11.73 cm. For VI, this corresponded to 15.90° of visual angle when the stimulus was at 42 cm from the observer (i.e., the diameter of the aperture plus two radii of the visual target). The stimuli followed a law of motion sinusoidal in acceleration defined by:

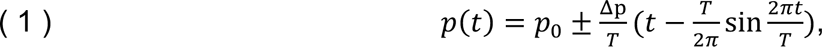

where *p* represents the vertical instantaneous position of the stimulus, *t* the instantaneous time elapsed from the beginning of motion, *p*_0_ the vertical initial position, and *T* the total duration of the stimulus. This law of motion was chosen based on previous research on vestibular motion perception^71^.

In each trial, we presented two consecutive stimuli moving vertically in opposite directions: a reference stimulus (R) and a comparison stimulus (C), the order of presentation of the two being randomly permuted, either R first (R/C) or C first (C/R). R had a fixed motion duration of *T* = 1.400 s, while C had 9 possible motion durations (ranging from *T* = 0.840 s to *T* = 1.960 s, in 0.140 s steps, corresponding to peak velocities of 27.93 cm/s for the fastest C and 11.97 cm/s for the slowest C).

In Figure 2, we show the stimulus trajectories for all R/C conditions in which the C stimulus was directed downwards. In all panels, from left to right, vertical dotted lines indicate the beginning and the end of the R stimulus and the start of the C stimulus. The position of the chair in world reference coordinates is shown in Figure 2a, d, g for VI, VE and VV, respectively. All possible durations of the C stimuli are color-coded lines from blue to red, from 0.840 s to 1.960 s, with the 1.400 s condition in black in the middle. In VI, the chair is shown motionless (Figure 2a). Figure 2b, e, h depict the visual target position (i.e., Target) in world reference coordinates for VI, VE and VV, respectively. Since the visual scene did not include Layer 1 in VE, the visual target position is absent in Figure 2e. The visual target was static in VV (Figure 2i). Finally, Figure 2c, f, i represent the visual stimuli resulting from the integration of visual and vestibular stimulation for VI, VE and VV, respectively. Notably, the visual stimuli were opposite between equivalent VI and VV conditions, while they were absent in VE conditions.

**Figure 2.**
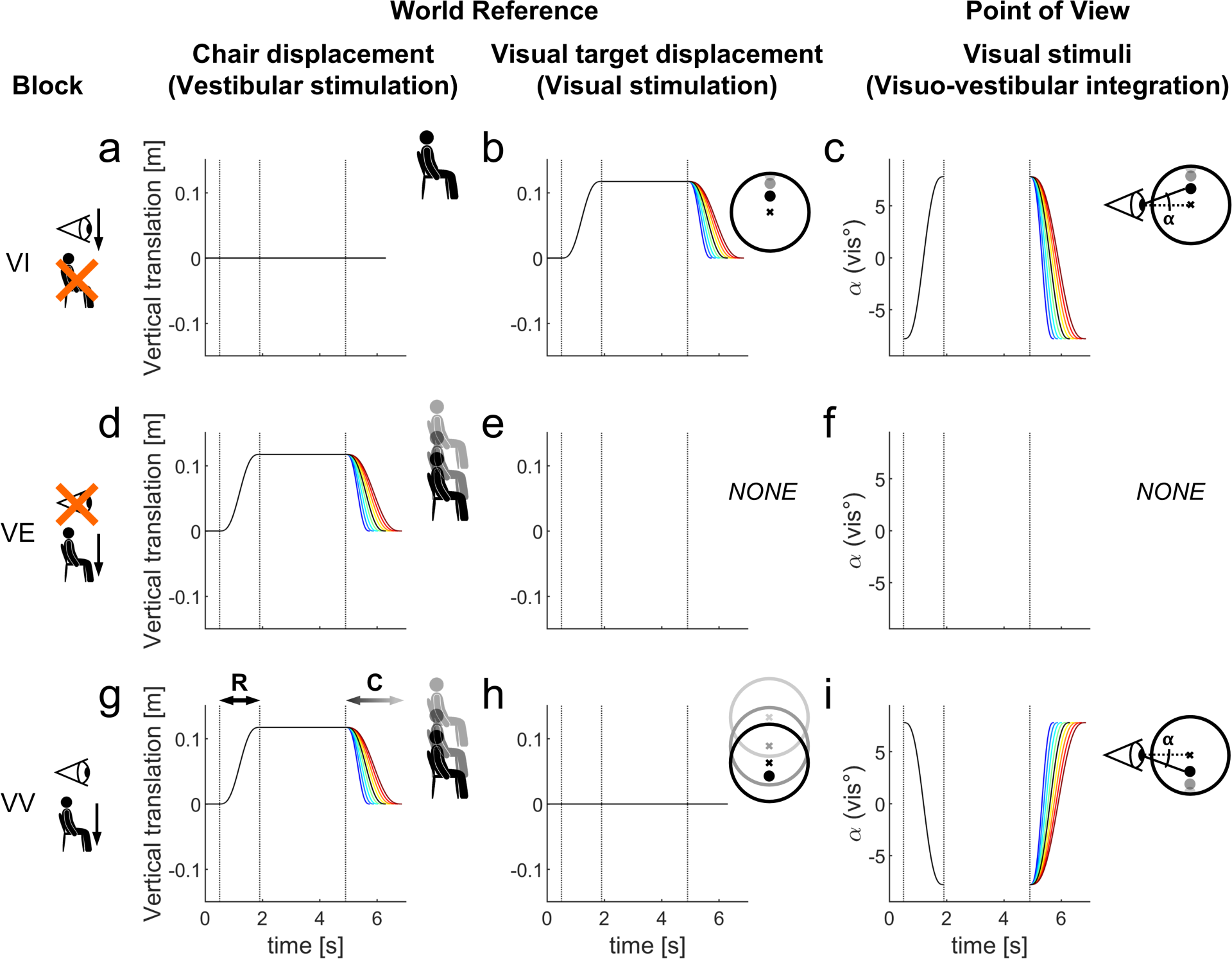
Vestibular and visual stimuli presented during the execution of the task in R/C conditions for all possible C durations (blue-dark red lines, in black the 1.400 s duration) in the blocks where C stimuli moved downwards (C direction Down); as depicted by the vertical arrows in block icons. VI (a-c), VE (d-f) and VV (g-i) sessions are hereby represented for their chair (a, d, g) and the visual target relative positions (b, e, h) in world reference. From the participant point of view, the visual stimuli visible through the VR-headset, resulting from visual (when present) and vestibular (when present) stimulations, are represented in panels c, f and I for VI, VE and VV sessions, respectively. In this representation the working area is 42 cm in front of the participant, as defined during the calibration phase (see text). In the sessions where the visual (VV) or vestibular (VI) stimulations were static, only a horizontal black line is depicted (as constant were the positions of the visual stimulus, panel h, or the chair, panel a, respectively). In VE no visual stimuli appeared on screen. Consequently, no positions are represented in e and f panels. In each panel from left to right, the three vertical dotted lines represent the beginning and the end of R and the beginning of C stimuli. C/R conditions were identical, changing only the order of presentation of C and R stimuli (not depicted). In C direction Up blocks, the C stimuli moved upwards, while the R stimuli moved downwards (not depicted).

### Motion platform control

Before the calibration phase, the MOOG motion platform was put on *engage*, at the center of its workspace. During the VI session, this position was maintained. In the VE and VV sessions, the motion of the chair was controlled in degrees of freedom at 1000 Hz by means of custom programs written in LabVIEW (2009, National Instruments, USA). All the details about the MOOG control are provided in La Scaleia et al.^70^.

### Protocol

Each session included two blocks identified by C direction: one block with C stimuli directed downwards and one block with C stimuli directed upwards. In VI, during the presentation of the two stimuli (R and C) in each trial, the visual target moved along the vertical diameter of the aperture, parallel to the dark-grey surface, following a trajectory defined by Equation 1. During the VE session, the chair moved vertically according to the same Equation 1. Finally, in the VV session, where both the visual stimulation and the vestibular stimulation were present, the chair moved as in the VE sessions, while the visual target was static. Therefore, in VV-Down (VV-Up), the chair moved downwards (upwards) for C stimuli. Moreover, in VV-Down (VV-Up), not only was the vestibular stimulus the same as that of the VE-Down (VE-Up) but, since the chair motion disclosed the static visual target, the resulting visual stimulation was comparable with that of VI-Up (VI-Down). See Figure 2 and the supplementary videos for further details.

### Procedure

Throughout each session, white noise was reproduced through the headphones at a volume high enough (75 dB) to ensure acoustic isolation of the participants. The beginning of each trial was signaled by a go-sound (81 dB pure tone at 500 Hz, 125 ms duration). Five hundred milliseconds after the onset of the sound, participants experienced the first stimulus (R or C). Then, the second stimulus (C or R) was presented after an interstimulus interval (ISI) of 3 seconds. This ISI should mitigate vestibular motion after-effects^72^ in VE and VV sessions, and should not involve major memory decays of the first stimulus in any of the sessions, including VI^43,73–75^. At the end of the second stimulus, a question appeared for 3 s on the screen at the center of the working area. The participants were asked to judge which one of the two stimuli had the shorter duration. The two possible answers were shown with a box on the left and an identical box on the right with “1” and “2” captioned on it, respectively. Responses were given by pressing a button on the Quest Pro left or right controller when the preferred answer was the first or the second stimulus. Visual feedback on the button that was pressed appeared on the screen by highlighting the corresponding box (“1” or “2”), and it stayed there until the question disappeared. In case of no response within the 3 s, no box was lit. No feedback about the correctness of the response was ever given. A new trial started 1 s after the disappearance of the question. In each trial of VI and VV sessions, participants were asked to keep fixation on the fixation cross. In each trial of VE sessions, participants were asked to fixate an imaginary target placed along the sightline and moving together with the subject.

The permutation of the levels of the two factors being manipulated – namely the 2 presentation orders of the stimuli within each trial (R/C or C/R) and the 9 possible durations of C stimuli – led to a total of 18 conditions, repeated 5 times each in a pseudorandom order. Thus, each experimental block consisted of 90 trials (18 conditions x 5 repetitions) for a duration of around 20 minutes. The two blocks of each session were separated by a 5-30 min break according to participants’ needs, while the three sessions were performed on different days (∼6 days apart). Both the sessions and the C direction order of presentation were counterbalanced across participants.

### Data Analysis

To assess the head stability, we measured in each experimental block the shift of the head in 3D relative to the calibration reference position, and the absolute value of the maximum shift of the head relative to the chair. To assess the maintenance of fixation, in each experimental block we computed average and standard deviation of the eyes shift from the reference position (i.e. the fixation cross in VI and VV sessions, or the imaginary target along the sightline estimated at the beginning of each trial in VE) on the horizontal and vertical axis.

We analyzed the psychometric results of the discrimination task by means of a two-level algorithm and a generalized linear mixed model (GLMM)^76^. For the two-level algorithm, first we fitted the valid responses of each participant separately for each block with a psychometric function (or general linear model):

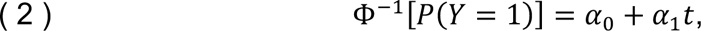

where *P*(*Y* = 1) is the probability of reporting that the comparison stimulus had a longer duration than the reference stimulus, Φ^–1^ is the probit link function, *α*_0_ and *α*_1_ are the intercept and slope of the general linear model, respectively, and *t* is C motion duration. To this end, we collapsed the responses for the order of presentation of the two stimuli (C/R and R/C). The point of subjective equality (PSE) and the just noticeable difference (JND) were computed from the equation ( 2 ) as in Moscatelli et al.^55^ for every block of each participant.

Finally, we also defined a GLMM to take into account the variability of the parameters estimates between subjects^76^. We fitted the data from different participants and blocks with a GLMM of the form:

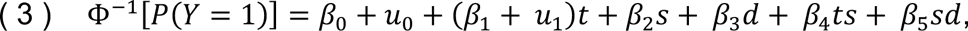

where the differences from the psychometric function are related to *t*, *s*, *d* accounting for the predictor variables, motion duration (*t*), session (*s*) and C direction (*d*). The parameters *β* and *u* represent fixed-effect and random-effects, respectively. We selected the model based on the lowest Bayesian Information Criterion (BIC), similarly to the procedures described in Moscatelli et al.^76^. Estimates on PSEs and JNDs were computed with a 95% Confidence Interval (95%CI) by means of the bootstrap method (2000 bootstrap sampled datasets).

In the Results section, we define *observed* PSE and JND values those derived from Equation 2, and *estimated* PSE and JND values those derived from Equation 3.

We used custom programs in MATLAB (R2023a, Mathworks, US) for data analysis.

### Statistics

Shapiro-Wilk test was used to assess the normality of the data distributions (alpha level = 0.05). For normally distributed data, we report mean values and 95%CI of the selected parameter over all participants (n = 20). Next, we defined a repeated measures ANOVA (RM-ANOVA) model, with session (3 levels: VI, VV and VE), direction of C stimuli (2 levels: Down and Up) and their interaction as within subject factors (Greenhouse-Geisser corrected). We performed a planned comparison by means of a pairwise comparison (t-tests, Bonferroni corrected). In particular, we excluded the comparison between VV and VI sessions within the same direction of C stimuli given that the visual stimulations in VV blocks were in opposite directions than those in corresponding VI blocks (see Figure 2 panels c and i). By contrast, we compared VE-Down (VE-Up) to VV-Down (VV-Up) and VI-Down (VV-Up) since the stimulations were in the same direction (see Figure 2 panels d and g and Figure 2 panels d and b for VE to VV and VE to VI, respectively). We performed additional statistical (t-test) analyses to directly compare VV blocks to the VE and VI blocks with similar vestibular and visual stimulations, respectively (i.e. VV-Down to VE-Down and VI-Up on the one side, VV-Up to VE-Up and VI-Down on the other side).

For not-normally distributed data, we report the median value and 95%CI and non-parametric statistics (Friedman test and Wilcoxon signed-rank test for planned comparisons, Bonferroni corrected).

We used custom programs in R (4.3.3, R Core Team) for statistical analyses.

## RESULTS

We first report the results about the compliance of the participants to the instruction of keeping the gaze fixed during all trials. Next, we report the psychophysical results of the discrimination of motion duration for the sessions with a visual target (VI), self-motion without an overt visual target (VE), and visuo-vestibular stimuli (VV).

### Head stability

Participants generally kept the head in a roughly constant position during the trials. At the start of the trial, the median shift of the head in 3D relative to the calibration reference position was -0.185 cm (95%CI [-0.582, 0.135], n = 60 (i.e. 3 sessions * 20 participants)), 0.050 cm (95%CI [-0.217, 0.180], n = 60), -0.756 cm (95%CI [-0.934, -0.585], n = 60) in x, y and z, respectively; and 0.162° (95%CI [-0.175, 0.368], n = 60), 1.520° (95%CI [0.825, 2.281], n = 60), -0.211° (95%CI [-0.599, 0.116], n = 60) in roll, pitch and yaw, respectively. At the start of the trial, no differences were observed in either displacements or rotations of the head between directions of C stimuli (Friedman test, all p > 0.179) and session (Friedman test, p > 0.086), except for roll (Friedman test, p = 0.043) whose changes were significantly smaller in VI than in VV (Wilcoxon test, p = 0.022, difference < 1°) and in VI than in VE (Wilcoxon test, p = 0.036, difference < 1°).

During each trial, the median absolute value of the maximum shift of the head relative to the chair position was 0.200 cm (95%CI [0.174, 0.216], n = 60), 0.177 cm (95%CI [0.163, 0.192], n = 60), 0.313 cm (95%CI [0.294, 0.329], n = 60) in x, y and z, respectively; and 0.286° (95%CI [0.261, 0.319], n = 60), 0.557° (95%CI [0.513, 0.629], n = 60), 0.264° (95%CI [0.244, 0.297], n = 60) in roll, pitch and yaw, respectively. The maximum head shift during trial presentation showed no significant differences in either displacements or rotations between directions of C stimuli (Friedman test, all p > 0.370). On the other side, it depended on the session (Friedman test, all p < 0.005, except for roll and yaw, p > 0.115), even though these differences were smaller than 2.5 mm and 0.1° for displacement and rotation, respectively. In particular, greater displacements and pitch rotations were observed in the sessions where the chair moved than in the visual session VI (Wilcoxon test, x direction: VI-VV p < 0.001 and VI-VE p = 0.132 after correction; y and z directions: all p < 0.001 after correction; pitch: all p < 0.004 after correction). Moreover, no significant differences were observed between VV and VE for displacements along the three axes and pitch rotations (Wilcoxon test, all p = 1.000 after correction).

### Fixation

Participants also maintained a good fixation during the trials. Eye position data showed that the median average shift from the reference position was 0.550° of visual angle (95%CI [0.464, 0.683], n = 60) and 0.873° of visual angle (95%CI [0.724, 0.963], n = 60) on the horizontal and vertical axis, respectively. The median standard deviation shift from the reference position was 0.329° of visual angle (95%CI [0.300, 0.379], n = 60) horizontally and 0.579° of visual angle (95%CI [0.493, 0.631], n = 60) vertically. Vertical eye average and standard deviation shifts were significantly larger than horizontal average and standard deviation eye shifts (Wilcoxon signed-rank test, p < 0.002).

No significant differences were observed in average or standard deviation shifts between directions of C stimuli (Friedman test, all p > 0.074) on both vertical and horizontal axis. Vertical average shifts also did not significantly differ across sessions (p = 0.387); however, vertical standard deviation shifts were significantly different across sessions (Friedman test, p = 0.004), though post-hoc tests revealed no significant pairwise differences (Wilcoxon tests, all p > 0.051 after correction, maximum difference < 0.28° of visual angle). Both horizontal average and standard deviation shifts were significantly different across sessions (Friedman test, all p < 0.005); however, post-hoc tests for horizontal average shifts revealed no significant differences (Wilcoxon tests, p > 0.079 after correction, maximum difference < 0.32° of visual angle). On the other side, horizontal standard deviation shifts exhibit significantly lower values in sessions with the fixation cross (i.e. VI and VV) than in VE (Wilcoxon test, all p < 0.001 after correction, maximum difference < 0.35° of visual angle). Noteworthy, previous studies have shown that horizontal eye positions standard deviations were smaller when fixating on a visible target compared to an imaginary target^77^.

### PSE values

As explained in the Methods, we computed the psychometric functions (Equation 2) for each condition and individual. They are plotted as Supplementary Figure 1, Supplementary Figure 2 and Supplementary Figure 3 for the VI, VE and VV condition, respectively. From each psychometric function, we derived the *observed* PSE (point of subjective equality). We found that these PSEs were normally distributed in all blocks (Shapiro-Wilk tests, all p > 0.050). The individual values and the box plots of the *observed* PSEs are depicted in Figure 3a. The mean values and 95%CI over all participants are reported in Table 1.

**Figure 3.**
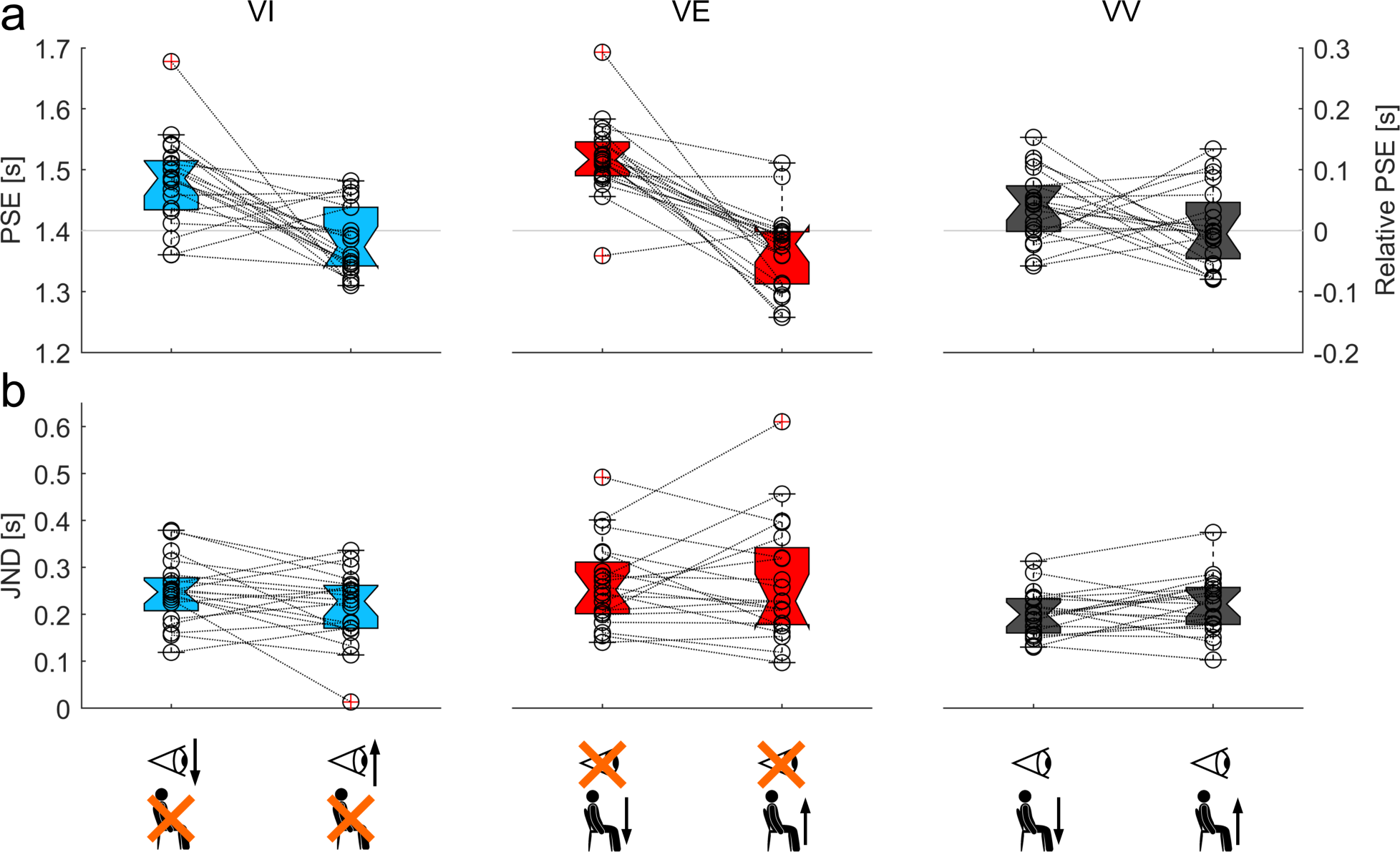
Boxplot for PSEs (a) and JNDs (b) obtained from all subjects in the two C directions of VI, VV and VE sessions. In light blue, red and dark grey data from VI, VE and VV sessions, respectively. Block icons in the same format as Figure 2. VV blocks are depicted according to their vestibular stimulation. Notches indicate 95%CI for the median. Data marked with red crosses indicate outliers (> ± 2.7 SD). In the a) panel, the continuous horizontal line represents the 1.400 s duration (i.e. the duration of R stimuli).

**Table 1.** Observed and estimated values (mean ± 95%CI) for PSE. Observed values are based on the individual psychometric functions defined on participants’ responses. Estimated values are the estimates of PSE estimated by the GLMM.

| Session | C direction | *Observed PSE [s]* |  |  | *Estimated PSE [s]* |  |  |
| --- | --- | --- | --- | --- | --- | --- | --- |
|  |  | Mean | -95%CI | +95%CI | Mean | -95%CI | +95%CI |
| VI | Down | 1.481 | 1.449 | 1.514 | 1.477 | 1.457 | 1.499 |
| VE |  | 1.520 | 1.492 | 1.547 | 1.512 | 1.489 | 1.536 |
| VV |  | 1.439 | 1.414 | 1.464 | 1.437 | 1.418 | 1.456 |
| VI | Up | 1.385 | 1.360 | 1.409 | 1.384 | 1.366 | 1.403 |
| VE |  | 1.367 | 1.339 | 1.369 | 1.370 | 1.349 | 1.391 |
| VV |  | 1.406 | 1.378 | 1.434 | 1.405 | 1.388 | 1.424 |

In the visual (VI) and vestibular (VE) sessions, we found a consistent trend across subjects for the values of PSE in the two motion directions (up or down) of the comparison (C) stimuli. In particular, we found that the mean PSEs over all participants were significantly (p < 0.050) larger than 1.400 s (i.e. the duration of R, the reference stimulus) when the C stimuli were directed downwards in both VI-Down and VE-Down blocks (right tailed one sample t-tests, all p < 0.001) (see Figure 3a). Conversely, the mean PSEs tended to be smaller than 1.400 s when the C stimuli were directed upwards in both VI-Up (left tailed one sample t-test, p = 0.121) and VE-Up (left tailed one sample t-test, p = 0.020). In other words, downward motion of the comparison stimuli was perceived as of the same duration as the upward motion of the reference stimulus only when it lasted longer than the reference, and vice versa for comparison stimuli moving upwards. At the individual level, the PSE for VI-Down was greater than the PSE for VI-Up in 16/20 participants, and the PSE for VE-Down was greater than the PSE for VE-Up in 19/20 participants.

The results were less consistent for the visuo-vestibular (VV) blocks (Fig. 3a). While the mean PSE over all participants was significantly (right tailed one sample t-test, p < 0.004) larger than the reference duration when the C stimuli were directed downwards (VV-Down), the mean PSE was not significantly different (left tailed one sample t-test, p = 0.655) from the reference when the C stimuli were directed upwards (VV-Up). Moreover, the trend of the PSE values as a function of the direction of motion of C stimuli was quite variable across participants. Thus, the PSE for VV-Down was greater than the PSE for VV-Up in 12/20 participants, and it was lower than the PSE for VV-Up in the remaining 8 participants.

Confirming the previous observations, the RM-ANOVA model for PSE showed a strong dependency of the *observed* PSEs on the C motion direction (F(1,19) = 53.387, p < 0.001 Greenhouse-Geisser corrected), and on the interaction between sessions (VI, VE, VV) and C motion direction (Up, Down) (F(2,38) = 6.263, p = 0.006, Greenhouse-Geisser corrected. See Figure 4 and Table 3). Planned pairwise comparisons highlighted a significant difference between the two directions of C stimuli in both VI and VE (p ≤ 0.001, Bonferroni corrected), with larger PSEs in VI-Down and VE-Down. No significant difference emerged between the two directions of C stimuli in VV, VV-Down and VV-Up (p = 0.160 after Bonferroni correction).

**Figure 4.**
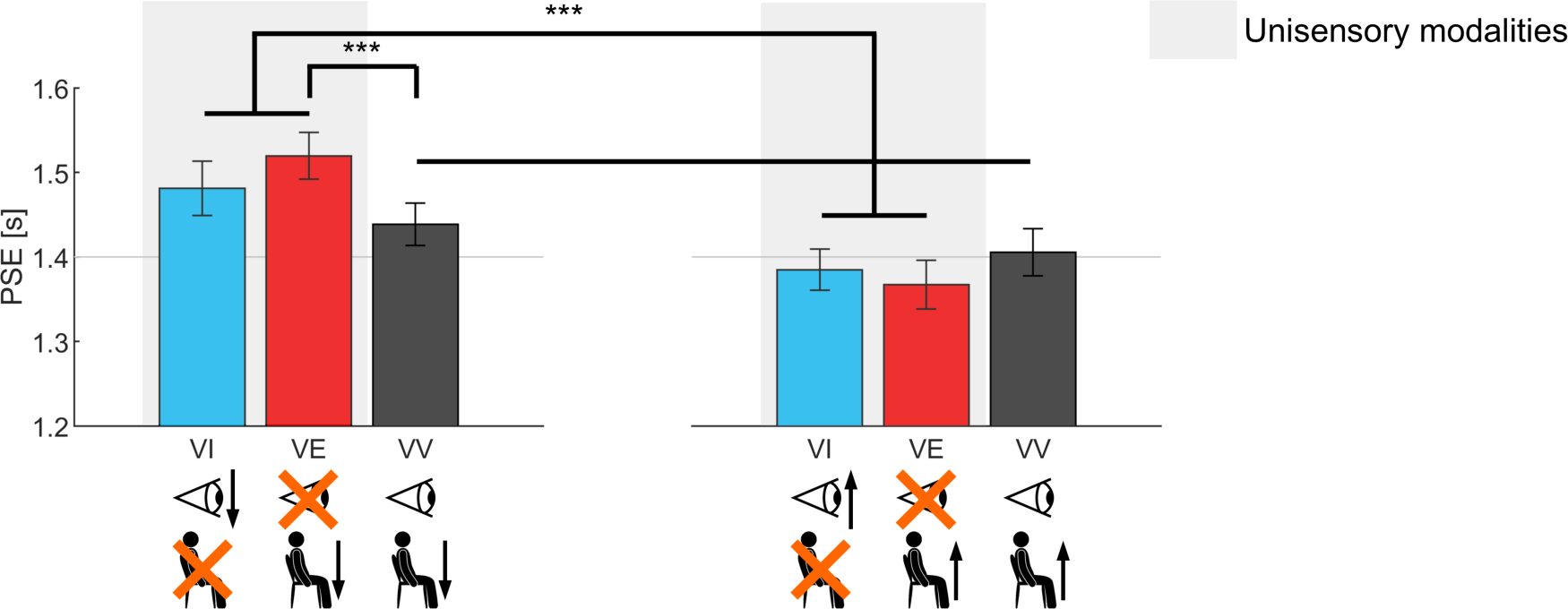
Bar plot for mean PSE values (error vertical line representing 95%CI) in C Down (left) and C Up (right), same icons and color code as Figure 3. RM-ANOVA for PSEs highlighted a significant effect of C direction and of the interaction between session and C direction. Planned pairwise comparisons highlighted that the latter effect is due to higher PSEs in VI-Down and VE-Down than in VI-Up and VE-Up, respectively, and to higher PSEs in VE-Down than in VV-Down. VV blocks are depicted according to the C direction of their vestibular stimulation. (**, p < 0.010. ***, p < 0.001).

**Table 2.** As for Table 1 for JND. Please note that estimated JNDs are identical across sessions of corresponding sensory modalities (see Methods and Table 4 for further details on the characterization of the GLMM).

| Session | C direction | *Observed JND [s]* |  |  | *Estimated JND [s]* |  |  |
| --- | --- | --- | --- | --- | --- | --- | --- |
|  |  | Mean | -95%CI | +95%CI | Mean | -95%CI | +95%CI |
| VI | Down | 0.248 | 0.218 | 0.278 | 0.239 | 0.218 | 0.260 |
| VE |  | 0.265 | 0.225 | 0.304 | 0.265 | 0.242 | 0.288 |
| VV |  | 0.201 | 0.180 | 0.223 | 0.217 | 0.200 | 0.235 |
| VI | Up | 0.215 | 0.181 | 0.248 | 0.239 | 0.218 | 0.260 |
| VE |  | 0.268 | 0.213 | 0.324 | 0.265 | 0.242 | 0.288 |
| VV |  | 0.219 | 0.192 | 0.246 | 0.217 | 0.200 | 0.235 |

**Table 3.** RM-ANOVA results for PSE and JND in the three sessions “s” (VI, VE and VV) and the two C directions “d” (Comparison -Down and Comparison -Up).

| factors | df1 | df2 | PSE |  | JND |  |
| --- | --- | --- | --- | --- | --- | --- |
|  |  |  | F | p | F | p |
| s | 2 | 38 | 1.658 | 0.204 | 5.789 | **0.006** |
| d | 1 | 19 | 53.387 | **< 0.001** | 0.188 | 0.670 |
| s * d | 2 | 38 | 6.263 | **0.006** | 1.915 | 0.165 |

Remarkably, the PSEs were not significantly different in VI and VE when the C motion direction was the same (p = 0.688 for the comparison VI-Down vs. VE-Down, and p = 0.184 for the comparison VI-Up vs. VE-Up, Bonferroni corrected). However, the PSE in Down was significantly shorter than in VE-Down (p < 0.001 after Bonferroni correction). The PSE in VV-Up was longer than in VE-Up, but the difference was not statistically significant (p = 0.160, after correction).

Since we found different PSE values when we applied a vestibular stimulation without overt visual stimuli in VE blocks and when we applied a vestibular stimulation in the presence of a static visual target in the VV blocks, we directly compared VV blocks with the VE and VI blocks with similar vestibular and visual stimulations, respectively (Figure 5a and Figure 5b). We found that the mean PSE value was significantly smaller in VI-Up than in VV-Down (paired t-test, p = 0.006 after correction), despite the visual target shifted upwards relative to the subjects’ fixation point in both cases. In addition, as previously noticed, the mean PSE value was significantly smaller in VV-Down than in VE-Down (paired t-test, p < 0.001 after correction), despite the subjects were displaced downwards in both cases (Figure 5a). Symmetrically, the mean PSE value was smaller in VV-Up than in VI-Down (paired t-test, p = 0.003 after correction), as it was the mean PSE in VE-Up relative to that in VV-Up (however, this difference was not significant, paired t-test p = 0.160 after correction) (Figure 5b).

**Figure 5.**
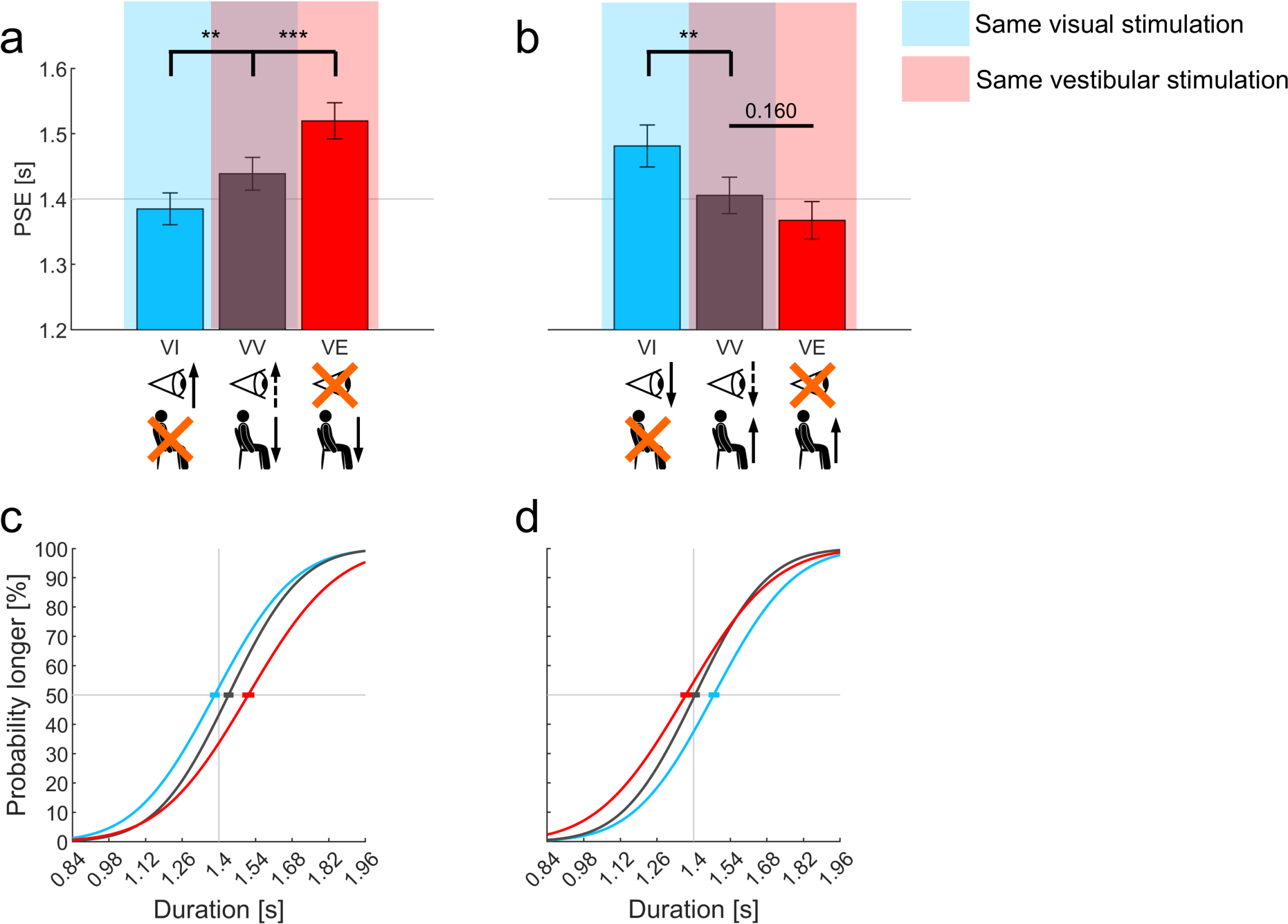
Bar plot for mean PSE values (error vertical line representing 95%CI) rearranged to match VV blocks with VI and VE blocks that correspond to equivalent visual and vestibular stimulations, respectively (panel a: VI-Up, VV-Down and VE-Down blocks; panel b: VI-Down, VV-Up and VE-Up blocks). Planned t-tests highlighted how PSEs from VV blocks fall in between values from unisensory conditions with correspondent visual and vestibular stimulations (**, p < 0.010. ***, p < 0.001). Continuous arrows represent the C direction in the same format as Figure 2. Dashed arrows represent the visual C direction resulting from chair motion. Panels c and d represent the psychometric functions defined from *estimated* PSE and JND (error horizontal line representing 95%CI on the PSEs) with the same rearrangement of panels a and b, respectively. Same color code as Figure 3.

To account for the variability of the results between subjects, we also performed a GLMM (Equation 3). The model parameters are reported in Table 4. The PSE values *estimated* by the GLMM are reported in Table 4, and are quite comparable to the *observed* PSE values. The psychometric functions derived from the GLMM are plotted for VV-Down (Figure 5c) and VV-Up (Figure 5d) so as to compare VV blocks with the VE and VI blocks with similar vestibular and visual stimulations, respectively.

**Table 4.** GLMM fixed effects parameters. t, motion duration; s, session; d, direction of C stimuli.

| Parameter | Estimate | Standard Error | z value | Pr(>\|z\|) |
| --- | --- | --- | --- | --- |
| (Intercept) | -5.683 | 0.272 | -20.891 | **< 0.001** |
| t | 3.758 | 0.190 | 19.714 | **< 0.001** |
| s VI | -0.473 | 0.228 | -2.080 | **0.038** |
| s VV | -0.908 | 0.237 | -3.828 | **< 0.001** |
| d Up | 0.533 | 0.056 | 9.531 | **< 0.001** |
| t * s VI | 0.411 | 0.149 | 2.764 | **0.006** |
| t * s VV | 0.829 | 0.157 | 5.268 | **< 0.001** |
| s VI * d Up | -0.147 | 0.080 | -1.835 | 0.067 |
| s VV * d Up | -0.388 | 0.082 | -4.748 | **< 0.001** |

### JND values

In addition to the *observed* PSE, the individual psychometric functions (Equation 2) yielded the value of *observed* JND (the just noticeable difference), which is inversely related to the precision of discrimination. We found that these JNDs were normally distributed in all blocks (Shapiro-Wilk tests, all p > 0.050). The individual values and the box plots of the *observed* JNDs are shown in Figure 3b. The mean values and 95% CI over all participants are reported in Table 1.

For all sessions (VI, VE, VV), the precision was similar between the Up and Down directions of C (paired t-tests, all p > 0.076) (Fig. 3b and Table 2). Moreover, the precision in VI and VE tended to be lower (i.e. higher JNDs) than in VV. Accordingly, RM-ANOVA of the *observed* JNDs showed a significant difference across sessions (F(2,38) = 5.789, p = 0.009, Greenhouse-Geisser corrected. See Table 3), mainly due to significantly higher JND values observed in VE than in VV (p = 0.021, Bonferroni corrected). JNDs were higher in VE than in VV in 11/20 and 18/20 subjects in the Down and Up conditions, respectively.

The results obtained with the GLMM (Equation 3) confirmed those reported above. The values of *estimated* JNDs were similar to the values of *observed* JNDs (see Table 2).

## DISCUSSION

In order to investigate whether the internal model of gravity affects visual, vestibular and visuo-vestibular perception of vertical motion duration, we designed a 2AFC task in which participants compared the durations of two consecutive stimuli moving in opposite vertical directions. The stimuli consisted of visual targets moving relative to the participant (VI), passive whole-body movements (VE), or their combination (VV) presented in three separate sessions. To reveal the presence of perceptual biases, we computed the point of subjective equality (PSE) of the psychometric functions with two different methods, finding very similar results.

Consistent with the prediction of the internal model of gravity, we found that downwardly directed motions of either the visual target (VI) or the participant (VE) were judged as lasting less (greater PSE) than upward motions of the same duration, and vice-versa for the opposite directions of motion. Moreover, the average PSEs over all participants for the corresponding visual and vestibular blocks (i.e. with the comparison stimuli moving up or down in both VI and VE blocks) were not significantly different, indicating that a similar perceptual bias related to gravity affects both sensory modalities. Our protocols involving both upward and downward motion in the same block (one direction for the comparison stimulus and the other one for the reference stimulus) did not allow separate estimates of precision (JND) for downward and upward motions.

The present results with visual stimuli are in agreement with previous findings by Moscatelli et al. (2019)^55^. Here, the observers were asked to compare the duration of downward versus upward motions, while in Moscatelli et al. the observers were asked to compare the speed of downward versus upward motions. The results also agree with the previous findings by Senot et al. (2005)^78^ that observers trigger hand movements to intercept a virtual target earlier when the target comes from above than when it comes from below, despite both target motions have the same durations and irrespective of their law of motion (1*g*, 0*g*, -1*g*). Therefore, the present and the previous studies reveal a bias consistent with the prior that downward motion is accelerated by gravity while upward motion is decelerated by gravity.

The present results with vestibular stimuli revealing an up-down bias identical to the visual bias are more novel. To our knowledge, previous studies focused on the discrimination of up-down directions in the dark, rather than motion duration (or speed) as in the present protocol. Some studies reported a higher precision (lower discrimination thresholds) for downward direction than for upward direction^45–48^, while other studies did not find a significant difference^49,63,64^. None of the above studies reported significant biases (asymmetries) between downward and upward directions in the upright posture (as in our participants), but their main focus was on the precision of discrimination. However, as mentioned in the Introduction, Kobel et al. (2021)^47^ found a main effect of motion relative to gravity due to a larger positive bias in earth-vertical motions than earth-horizontal motions for 2-Hz displacements. Since no significant bias was found for 1-Hz displacements where vestibular cues dominate over somesthetic cues^65,66^, the bias at 2-Hz could have resulted from up-down asymmetries in tactile cues^46,47^. However, the general conclusion of Kobel’s study is in agreement with the hypothesis that internal models of gravity influence the perception of vertical translations, since they found that earth-vertical thresholds, where the translation stimulus must be disambiguated from the colinear gravitational acceleration, were significantly higher than earth-horizontal thresholds, where the translation is independent of (i.e., perpendicular to) gravitational acceleration^47^.

While we cannot rule out the influence of somesthetic cues in the responses to up-down displacements of the participants, we notice that the present stimuli had frequencies between about 0.5 Hz and 1.2 Hz, thus near the range where vestibular cues are known to dominate over somesthetic cues^65,66^. Bruschetta et al. (2021)^79^ have specifically dissected the role of somesthetic versus vestibular cues in the perception of longitudinal and lateral motions in the dark. They found that somesthetic cues play a significant role for strong accelerations, while our stimuli were smooth ramp and hold waveforms. Moreover, it is known that the perception of earth-vertical translations –such as the present ones– is especially compromised by complete bilateral vestibular ablation, as shown by very high thresholds of vertical direction discrimination in these patients^65,66,80^. Nevertheless, even if the perceptual responses during VE (and VV) sessions included some non-vestibular sensory (e.g., somesthetic and visceral) contributions in addition to vestibular contributions, this would not change the conclusion that the judgment of duration of vertical whole-body motion is biased by the prior assumption about the effects of gravity.

We do not know where in the brain the up-down visual and (primarily) vestibular biases arise. Directional anisotropies in neural responses to either visual^81,82^ or vestibular^83^ stimuli have been described, mainly related to the directional preference for stimuli oriented along the cardinal axes relative to oblique stimuli. However, we are not aware of neural responses that can be reconciled with the downward bias that we described. For instance, in the monkey the responses of primary vestibular afferents preferentially excited by upward translations are similar to the responses of afferents preferentially excited by downward translations^84^. Also the responses of Medial Superior Temporal (MST) area neurons to upward and downward body translations are similar^85^. By the same token, multiple measures of Middle Temporal (MT) visual area neuronal responses do not provide evidence of a directional anisotropy to visual motion^86^.

Downward biases may result from processing at neural stages downstream of MT/MST, in particular at the level of the parieto-insular vestibular cortex where the internal model of gravity effects has been shown to be encoded in fMRI^3,21,87–89^ and TMS studies^23,24^. Neural populations in the temporo-parietal junction, posterior insula, retro-insula and OP2 in the parietal operculum are selectively engaged by vertical visual motion of objects and self-motion coherent with gravity, as well as by vestibular stimuli^3,21,22,90,91^. Patients with lesions of the temporo-parietal junction and peri-insular regions show specific deficits in the perceptual estimates of passive motion durations in the dark^92^, as well as deficits in the processing of visual targets accelerated by gravity^93^.

On the other hand, we did not expect systematic biases in the visual-vestibular sessions (VV), given that the vestibular stimulus was in the opposite direction of the visual stimulus, with the result that the visual and vestibular bias should cancel out. In fact, on average there was no significant difference in the bias (PSE) between the block with downward motion of the participant and that with upward motion. However, this lack of significant difference of the average values did not seem to depend on the individual values of the visual and vestibular biases cancelling out. It rather depended on a large inter-subject variability of PSE values: in about half of the participants, the PSE for downward self-motion was greater than that for upward self-motion, but in the other half of the participants the opposite was true. This result suggests that each subject weighed relatively more one cue (visual or vestibular) than the other one, since visual motion and vestibular motion were in opposite directions. Inter-subject variability might depend on the different way each participant resolved the potential ambiguity inherent in our VV protocol. While the participants underwent whole-body vertical motion, the visual target was static throughout and it only appeared to move in the opposite direction of the body due to body motion.

Nevertheless, some evidence for visual-vestibular fusion in VV sessions is provided by the significantly higher average precision (lower JND) than in the vestibular sessions (VE).

## Conclusion

We confirmed and extended previous findings that downward visual targets are perceived as faster (shorter duration) than upward targets. We further showed that the same bias occurs with whole-body motion. When visual and vestibular cues were combined together, on average the downward bias disappeared. Overall, the results suggest that visual and vestibular modalities of motion perception share the same prior assumption about the effects of gravity, putatively encoded in the parieto-insular vestibular cortex.

## ACKNOWLEDGEMENTS

This study was supported by grants from the Italian Space Agency (I/006/06/0), INAIL (BRIC 2022 LABORIUS), Italian University Ministry (PRIN 2020EM9A8X, 2020RB4N9, 2022T9YJXT, 2022YXLNR7, #NEXTGENERATIONEU NGEU National Recovery and Resilience Plan NRRP, project MNESYS PE0000006 – A Multiscale integrated approach to the study of the nervous system in health and disease DN. 1553 11.10.2022) and Space It Up project funded by the Italian Space Agency and the Ministry of University and Research - Contract No. 2024-5-E.0 - CUP No. I53D24000060005. The authors thank Greta Dimasi for help with the setup.

## AUTHOR CONTRIBUTIONS

SDM and BLS conceived the study, researched and analyzed data, and wrote the first draft of the manuscript. AFA contributed to the characterization of the setup. FL and MZ contributed to discussions and successive drafts of the manuscript. MZ contributed resources. All authors approved the final version of the manuscript.

## DATA AVAILABILITY

Data analyzed in this study are included in the published article and its online supplementary file. Additional data are available from the corresponding authors upon reasonable request.

## ADDITIONAL INFORMATION

The authors declare that they have no competing interests related to this work.

## SUPPORTING INFORMATION

Additional supporting information can be found online in the Supporting Information section at the end of this article, hosted online.

## SUPPLEMENTARY FIGURES

**Supplementary Figure 1.**
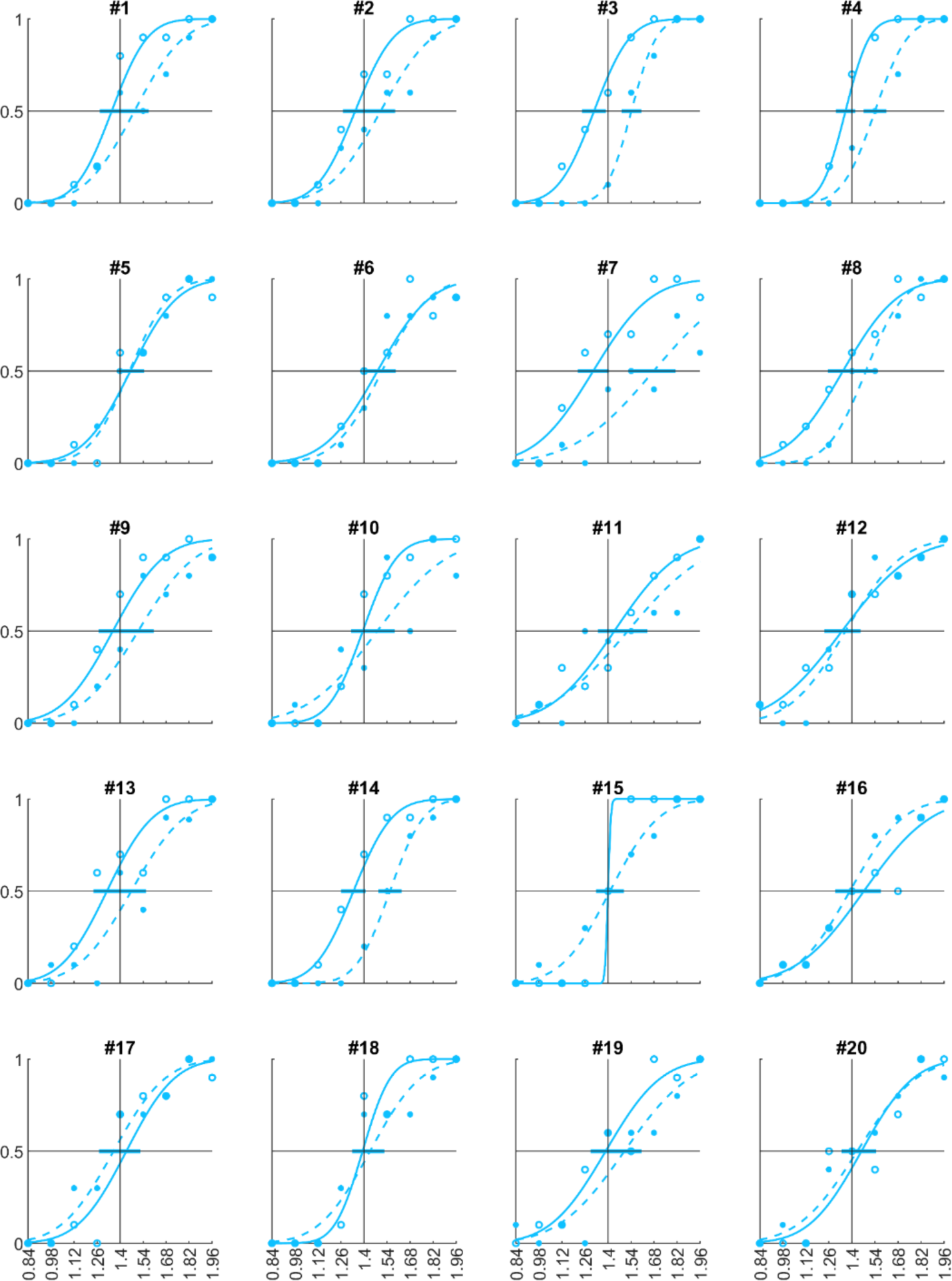
Psychometric functions from the two visual blocks (VI) of each participant (error horizontal line representing 95%CI on the PSEs). PSEs and JNDs extracted from these curves are those reported in the boxplots of Figure 3. In filled circles and continuous line data from Up, in open circles and dashed line data from Down.

**Supplementary Figure 2.**
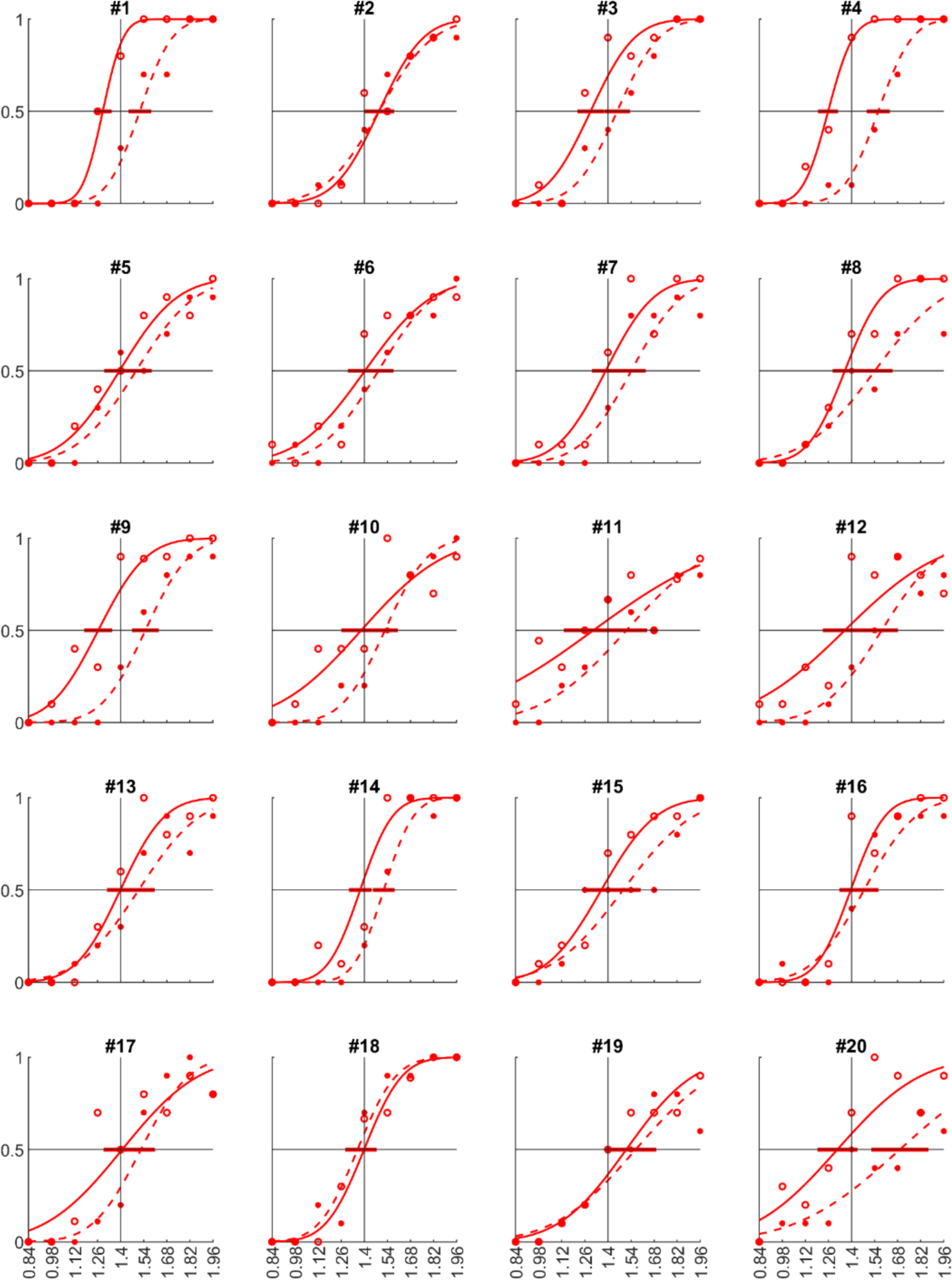
Psychometric functions from the two vestibular blocks (VE) of each participant (error horizontal line representing 95%CI on the PSEs). PSEs and JNDs extracted from these curves are those reported in the boxplots of Figure 3. Same layout as Supplementary Figure 1.

**Supplementary Figure 3.**
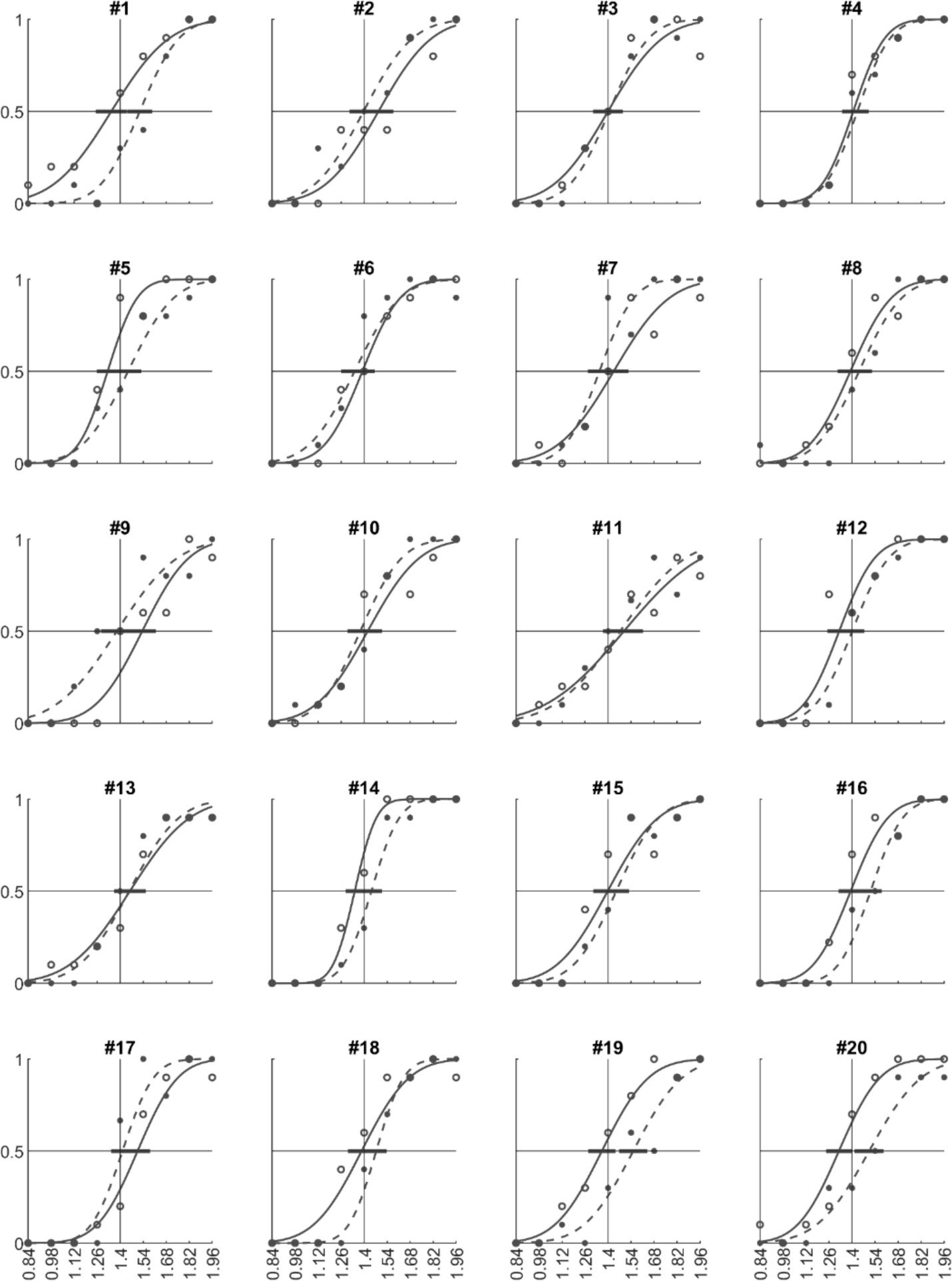
Psychometric functions from the two visuo-vestibular blocks (VV) of each participant (error horizontal line representing 95%CI on the PSEs). PSEs and JNDs extracted from these curves are those reported in the boxplots of Figure 3. Same layout as Supplementary Figure 1.

## SUPPLEMENTARY VIDEOS

Supplementary Video 1. One R/C 840 ms trial in VI-Down (see also Figure 2a-c, black/dark-blue trajectories). On the left side, the world reference view of the vestibular and visual stimulation. The subject sat on a chair with their unrestrained head facing forward while wearing a VR-headset system. In the 3D virtual scenario displayed in it, the position of the working area (Figure 1) was defined during the initial calibration phase by placing the fixation cross 42 cm in front on their antero-posterior axis. From that instant, the position of the aperture was matched to that of the chair. In the world reference visual stimulation column, the visual target can be seen to appear from behind the aperture. On the right side, the point of view of the participant when the head was facing forward.

Supplementary Video 2. One trial R/C 840 ms in VE-Down (see also Figure 2d-f, black/dark-blue trajectories). Same layout as in Supplementary Video 1.

Supplementary Video 3. One trial R/C 840 ms in VV-Down (see also Figure 2g-i, black/dark-blue trajectories). Same layout as in Supplementary Video 1. Noteworthy, during VV blocks the position of the visual target was static. Chair motion temporarily disclosed it, leading to a visual stimulation that, from subject’s point of view, was opposite in direction to that in Supplementary Video 1.

